# Global Gene Expression Divergence in Spontaneous Mutation Accumulation Lines of *Caenorhabditis elegans* under Varying Efficiency of Selection

**DOI:** 10.1101/2021.05.03.442498

**Authors:** Thaddeus C. Deiss, Anke Konrad, Ulfar Bergthorsson, Vaishali Katju

## Abstract

To ascertain the effect of relaxation of selection on global gene expression, *Caenorhabditis elegans* mutation accumulation (MA) lines were propagated under varying degrees of efficiency of selection determined by their different population sizes (*N* = 1, 10 and 100). Both the mutational variance (*V*_*m*_), and the residual variance (*V*_*r*_) were greatest in MA lines with the lowest efficiency of natural selection. The results suggest that gene expression is under strong balancing selection. Furthermore, mutations resulting in increased transcriptional noise or sensitivity to microenvironmental variation accumulate most under extreme genetic drift. In contrast, the *V*_*m*_/*V*_*r*_ ratio was lowest in the *N* =1 lines. Chromatin domains associated with broad gene silencing and active transcription exhibited the greatest and the smallest increase in transcriptional variation, respectively. Furthermore, the preponderance of overexpressed genes was especially pronounced in mitochondrial respiration, stress response, and immune system pathways, especially in low fitness *N* = 1 lines.

## INTRODUCTION

Experimental investigations into the rate of spontaneous mutations, along with their molecular and phenotypic consequences, are central to the study of evolution and biology, with important implications for human health including disease susceptibility and inherited genetic disorders. Regulation of gene expression has long been understood to play a considerable role in evolution and most phenotypic evolution, both intraspecific and interspecific, is likely due to changes in gene expression rather than changes in protein structure (Britten and Davidson 1969; King and Wilson 1975; Stern 1998; Wittkopp *et al*. 2003; Wray *et al*. 2003; Abzhanov *et al*. 2004; Fay *et al*. 2004; Shapiro *et al*. 2004; Gompel *et al*. 2005; Gilad *et al*. 2006; Fay and Wittkopp 2008; Loehlin *et al*. 2019).

The phenotypic consequences of spontaneous mutations in their broadest sense, including behavioral, biochemical and physiological traits, are realized and constrained by the rules of gene expression and development. Characterizing patterns of gene expression divergence in natural and experimental populations is therefore an important step towards understanding how mutations and epimutations exert their influence on the evolution of gene expression. Furthermore, questions regarding the relative importance of natural selection versus genetic drift in shaping the evolution of expression require knowledge about the effects of spontaneous mutations on gene expression. The influence of these two major evolutionary forces, drift and selection, can be investigated by comparing populations or experimental laboratory lines under minimal selection (strong genetic drift) with their counterparts that are subject to strong intensity of selection (e.g. natural populations) (Denver *et al*. 2005; Rifkin *et al*. 2005; McGuigan *et al*. 2014b; Huang *et al*. 2016). Moreover, just as genes that differ markedly in their rates of sequence evolution have provided evidence for selective constraints and positive selection, genes may also exhibit differential capacity to evolve at the transcriptional level (Rifkin *et al*. 2005; Landry *et al*. 2007). In a related vein, gene expression divergence may also vary across developmental stages, exhibiting highly constrained patterns in some stages (developmental constraints) relative to others (Zalts and Yanai 2017).

Additional questions of interest relate to the nature of genetic change driving divergence of gene expression. Given the considerable variation in genome organization of different groups of organisms, how might a species’ prevailing genomic and genetic architecture impinge on the evolution of its transcriptome? The genomes of eukaryotic species are highly variable in size, chromosomal organization, and chromatin state, and can comprise large expanses of repetitive, gene-poor regions of low complexity as well as a high incidence of selfish genetic elements. Additionally, there exists considerable variation in recombination frequency which, in conjunction with selection, can further influence the patterns of nucleotide variation, and in turn effect differential divergence at the transcriptional level. Investigating how these different properties of spontaneous mutations and genome architecture influence the evolution of gene expression is of fundamental importance.

In order to determine the effective role of mutation in shaping expression profiles, many studies opt to negate the force of selection acting constitutively within natural populations. Mutation accumulation (MA henceforth) is a commonly implemented experimental design to isolate the effect of mutation from selective pressures (Halligan and Keightley 2009; Katju and Bergthorsson 2019). In an MA experiment, multiple replicate lines derived from an inbred ancestral stock population are allowed to evolve independently of one another under conditions of extreme bottlenecking each generation. The repeated bottlenecks severely diminish the efficacy of natural selection, promoting evolutionary divergence due to the accumulation of deleterious mutations by random genetic drift. While spontaneous MA experiments provide a powerful framework to investigate divergence in global transcription profiles due to accumulated genetic changes in the near absence of selection, only a limited set have been analyzed at the transcriptional level. These studies have determined global mutational variance (*V*_*m*_) in a few eukaryotic species comprising nematodes (Denver *et al*. 2005), flies (Rifkin *et al*. 2005; McGuigan *et al*. 2014a, 2014b; Huang *et al*. 2016), and yeast (Landry *et al*. 2007). More recent studies have focused on the transcriptional profiles of copy-number variants (CNVs) and transposable elements in *Caenorhabditis elegans* following MA (Konrad *et al*. 2018; Bergthorsson *et al*. 2020). Some initial conclusions from these studies suggest that gene expression divergence is under strong stabilizing selection. But despite strong selection, there is also a correlation between transcriptional effects of spontaneous mutations and divergence between closely-related species, suggesting that mutational input exerts a significant influence on the patterns of gene expression divergence (Rifkin *et al*. 2005).

Although MA experiments are canonically designed to negate the effect of selection by utilizing experimental lines repeatedly subjected to extreme population bottlenecks, this study aimed to identify trends in expression variability and divergence within lines subjected to varying efficacy of selection (Konrad *et al*. 2017, 2018, 2019). Experimental MA lines were maintained at three population sizes of *N* = 1, 10, and 100 individuals per generation as outlined in Katju *et al*. (2015, 2018), thereby enabling comparison of transcript abundance between populations evolved under the near absence of selection (*N* = 1) versus those subject to increasing efficacy of selection (*N* =10, and 100). A comparison of the mutational and residual variance between these different population size treatments can be used to investigate the effects of selection and drift on both the mutational heritability and plasticity in gene expression.

## METHODS

### Experimental study system and mutation accumulation design

We conducted a long-term spontaneous MA experiment in *C. elegans* comprising 35 populations maintained in parallel at varying population size treatments of *N* = 1, 10, and 100 hermaphrodites (*N*_e_= 1, 5, 50 individuals, respectively) per generation over four and a half years and spanning 409 MA generations (Katju *et al*. 2015, 2018). All experimental lines of the MA experiment were established from the descendants of a single wild-type Bristol (N2) hermaphrodite originally isolated as a virgin L4 larva (**Supplemental Figure S1A**) with excess animals cryogenically preserved at –86°C as ancestral controls. The MA experiment was conducted under benign laboratory conditions. Additional details on methodology are provided in Katju *et al*. (2015). Stocks of the MA lines were cryogenically preserved every 50–100 MA generations during the course of the experiment.

### Theoretical underpinnings

Mutational fitness effects can range continuously from lethal to deleterious to neutral to beneficial. Mutations in small populations are governed by genetic drift, with beneficial mutations lost and detrimental mutations fixed randomly within the population. However, as population size increases, natural selection steadily supplants genetic drift in dictating which mutations are lost versus fixed. Hence, the fate of a novel mutation, be it loss or fixation, is dependent upon both the selection coefficient (*s*) and effective population size (*N*_*e*_). The mutational dynamics for sexually reproducing diploids has been shown to be dominated by random drift when |*s*| << 1/2*N*_*e*_ (Kimura 1962). In contrast the dynamics for mutations with |*s*| >> 1/2*N*_*e*_ are dictated by natural selection. Complete inbreeding of obligate self-fertilizing hermaphrodites, such as *C. elegans*, results in a 50% reduction in *N*_*e*_ relative to the census population size (*N*) (Crow and Kimura 1970; Pollak 1987; Charlesworth 2009). This means that the effective population sizes for our experimental lines of *N* = 1, 10 and 100 individual(s) correspond to *N*_*e*_ = 1, 5 and 50 individuals respectively. The fate of mutations in our MA lines of *N* = 1, 10 and 100 is expected to be governed by genetic drift when selection coefficients are less than 0.5, 0.1 and 0.01 respectively. The demarcation between the behavior of mutations with 2*N*_*e*_*s* < 1, which should be dominated by genetic drift and 2*N*_*e*_*s* > 1, which are under greater influence of natural selection, is not sharply defined and some mutations with a larger fitness cost than 1/2*N*_*e*_ could be fixed by genetic drift. Nonetheless, the differences in populations size in these MA experiments alters the relative importance of genetic drift versus natural selection in the fixation or loss of mutations, with genetic drift having the greatest influence in *N* = 1 lines and diminishing in strength with increasing population size.

### RNA extraction, library preparation and RNA-Seq

One individual was isolated from each of 17 *N* = 1 lines, while two and three individuals were isolated from MA lines of size *N* = 10 (10 lines), and *N* = 100 (five lines), respectively. Given that the *N* =1 MA lines are expected to be genetically identical and homozygous due to self-fertilization for several hundred generations and extreme bottleneck size, we only isolated a single individual per *N* = 1 MA line for RNAsequencing. MA lines maintained at larger population sizes were expected to harbor greater standing genetic variation. Hence, we isolated two and three randomly picked individuals from each *N* = 10 line and *N* =100 line, respectively. Additionally, five randomly picked individuals were isolated from the pre-MA ancestral control. The isolated individuals were sequestered on to NGM plates containing OP50 lawns and kept at 20°C. For each of these 57 individuals, three offspring at the L4 larval stage were isolated from the F_1_ generation to serve as biological replicates in the expression analysis (**Supplemental Figure S1B**), yielding a total of 171 samples for tissue collection and RNA sequencing at the L1 larval stage. All 171 sample populations were maintained for a few generations until enough worms were available for RNA extraction. Synchronized populations of L1 larvae were generated through the collection of gravid eggs from adults using a standard bleaching protocol and plated on unseeded NGM plates. Eggs from the synchronized populations were allowed 12 hours to hatch and L1 larvae were collected for total RNA extraction. Total RNA isolation was performed using the Qiagen RNeasy kit (Cat no. 74106) according to manufacturer’s protocol with addition of a glass bead beating step to aid in nematode lysis. Quantification and assessment of RNA integrity was determined via Qubit 3.0 Fluorometer, HS RNA assay (Cat no. Q32852), and Agilent TapeStation, HS RNA assay (Cat no. 5067-5580), respectively. Illumina libraries were constructed from 0.5 µg of total RNA with the Illumina Truseq stranded mRNA kit and adapter indexes (Cat no. 20020594/20019892) according to manufacturer protocols. Libraries were pooled at a concentration of 5 nM for paired-end sequencing on the Illumina HiSeq4000 platform (2 × 150bp) at the Texas A&M Genomics and Bioinformatics Service Center or Illumina Novaseq6000 (2 × 150bp) at the North Texas Genome Center at the University of Texas at Arlington. Sequence data are deposited to the NCBI database with the accession number for the Bioproject PRJNA448413.

### Genomic Mapping and Analysis of Transcript Abundance

Sequence mapping and transcript quantification were performed with the STAR alignment software (Dobin *et al*. 2013) using default settings and the *C. elegans* WS274 genome sequence and annotation files as references for mapping and transcript quantification, respectively. Gene specific read counts for each biological replicate were used to construct a raw matrix for all downstream analyses. Transcripts per million (TPM) were calculated gene-wise, using the longest transcript isoform for alternatively spliced genes, for each sample using the following equation:

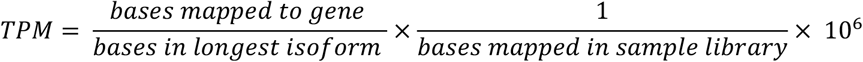

Genes with less than an average three TPM cutoff across all samples were excluded in all subsequent analyses. The filtered dataset contained 11,686 genes on which all downstream analyses were performed. Unless otherwise specified, the ratio of the MA line to the ancestral TPM values were log_2_-transformed prior to all downstream analyses to facilitates comparisons to previously reported microarray results.

Differential expression analyses were performed on TPM values using a moderated T-test (Smyth 2005) in the “limma” R package (Richie *et al*. 2015) with a design matrix built to compare the individual biological replicates for MA lines to the ancestral control biological replicates. The resulting *p-values* were adjusted using the Benjamini-Hochberg FDR correction for false discovery rate in multiple comparison testing. Differential expression analysis results were filtered with thresholds for adjusted *p-value* (adjusted *p* ≤ 0.01) and magnitude (20% change over ancestral N2 control, log_2_-fold change = 0.26). In summation the differential expression analysis returned comparisons for 3 × 17 samples from the *N* = 1 populations, 3 × 20 for *N* = 10, and 3 × 15 for *N* = 100 to the ancestral control.

Gene Ontology (GO) enrichment analysis was performed on the resulting differential expression gene lists with the “clusterProfiler” package (Yu *et al*. 2012) in R. The background gene list used for significance testing was composed of genes that were found to be differentially expressed between any MA sample and the ancestor regardless of population size. The minimum number of background genes per category was 50 while the number of differentially expressed genes per experimental category was set at five. A Benjamini-Hochberg adjusted *p-value* of 0.05 was used as a significance cutoff for GO category enrichment. A Wilcoxon-Signed Rank test was employed to determine significant differences in the number of underexpressed to overexpressed genes for all experimental populations. Population-specific differences in the distributions of overexpressed, underexpressed, and overexpressed:underexpressed genes were determined using Kruskal-Wallis tests.

### Estimation of the Mutational and Residual Variance

Estimates of residual variance (*V*_*r*_) and mutational variance (*V*_*m*_) were calculated in R (R Core Team 2014) using the model below for each gene allowing for population-specific across-line variance and line-specific residual variances:

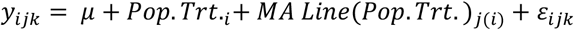

where *y*_*ijk*_ is the gene-wise log_2_-ratio of the MA sample to average ancestral value for each MA sample *k*, of each MA line *j*, evolved under population size treatment *i*. Mutational variance (*V*_*m*_) was calculated as the variance across experimental MA lines for each population size treatment divided by 2× the average number of generations per line for each population. This calculation yielded a single *V*_*m*_ value per gene for each population size treatment (*N* = 1, 10 and 100) which was then used to perform KruskalWallis tests for differences in the distribution of *V*_*m*_ between the three population size treatments.

The residual variance (*V*_*r*_) was calculated gene-wise for the three biological replicates of each genotype via two different methods depending on the downstream statistical comparisons. The first calculation used raw TPM values to compare *V*_*r*_ between MA lines and the ancestral control. The second approach used the log_2_-ratios of MA line replicates to ancestral control to determine *V*_*r*_ in the same manner as was used for calculating *V*_*m*_. Both of these calculations yielded gene-specific *V*_*r*_ estimates for each MA lines in the study. A Kruskal-Wallis test was used to investigate differences in the distribution of *V*_*r*_ between the three MA treatments (*N* = 1, 10, and 100) and the ancestral control). Following a significant Kruskal-Wallis test, a Dunn’s Multiple Comparison Test test with a Benjamin-Hochberg stepwise adjustment was used to conduct multiple pairwise comparisons for median difference. Each of the plate used for extracting RNA contained several thousand nematodes. Because the number of worms per RNA extraction for each biological replicate was unknown, we did not estimate the mutational heritability 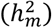. Under the assumption that there were no systematic differences in sample size between MA lines, we calculated the *V*_*m*_*/V*_*r*_ ratio for different population size treatments in lieu of 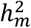.

### Histone Modification Datasets and Analysis

Genes were additionally assigned to one or more histone modification categories based on a previous *C. elegans* ChIP-seq expression study (Liu *et al*. 2011). Publicly available ChIP-Seq datasets were downloaded from the modMine repository (http://intermine.modencode.org) (Contrino *et al*. 2012). **(Supplemental Table S1)**. Datasets lacking pre-analyzed coordinates were downloaded in wiggle (WIG) format for analysis of peaks using the MACS2 software package (Zhang *et al*. 2008). A minimum interval of 1000bp was used to determine histone associated genomic tracks with the default peak cutoff suggested by the cutoff-analysis tool. Since our samples were L1 larvae and publicly available datasets were mostly composed of early embryo or L3 larvae, we only assigned genes to a category if there was a consensus amongst all ChIPSeq datasets for a given modification. For example, genes were only assigned to the *H3K27me1* category if they were found within a positive track for all three publicly available CHiP-Seq datasets.

### Statistical Analyses

All calculations and statistical tests were performed in R with graphs produced using the ggplot2 package (Wickham 2016).

## RESULTS

We employed a spontaneous MA experiment design in *C. elegans* comprising three population size treatments of *N* =1, 10, 100 individuals all descended from a single N2 worm ancestor (Katju *et al*. 2015, 2018). The MA lines of differing population sizes were subjected to bottlenecks for >400 consecutive generations, allowing us to jointly assess the genome-wide, molecular consequences of different mutation classes (in both the mitochondrial and nuclear genomes) under conditions of neutrality and with increasing intensity of selection (Konrad *et al*. 2017, 2018, 2019). In two preceding studies, we have assessed the transcriptional consequences of copy-number variants (gene duplications and deletions) and transposable elements (TEs) (Konrad *et al*. 2018; Bergthorsson *et al*. 2020). In both these studies, the *N* =1 lines that were subject to the greatest intensity of genetic drift displayed the greatest increase in transcriptional activity in gene duplications and TEs relative to the ancestral control. In this study, we further extend the transcriptional analyses of these evolved experimental lines to investigate the divergence in global transcription profiles due to the input of other classes of new mutational variants (excluding gene copy-number changes and TEs), both under minimal influence of purifying selection (*N* =1 lines) and with incrementally increasing efficacy of selection (*N* = 10 and 100 lines).

### Greater mutational and residual variance at small population size

We have previously reported that gene copy-number variants (CNVs) are more common in the *N* = 1 MA line, and that the increase in transcript abundance associated with gene duplications is also greater in the *N* = 1 MA lines compared to the *N* = 10 and *N* = 100 MA lines (Konrad *et al*. 2018). In addition, TE transcription exhibits a negative relationship with population size in MA lines (Bergthorsson *et al*. 2020). The results presented here were calculated after filtering out genes that were known to be duplicated or deleted (CNVs), as well as annotated transposable elements (TEs). The mutational variance, *V*_*m*_, is a fundamental parameter in models of quantitative traits, and is defined as the rate of increase in genetic variation per generation due to mutation. *V*_*m*_ was estimated as the increase in the variance between MA lines within a particular population bottleneck size standardized by 2× the average number of generations. The median *V*_*m*_ values for transcript abundance after removing CNVs were 1.11, 0.89, and 0.85 ×10^−4^ per gene for the *N* = 1, 10 and 100 lines, respectively (**Figure 1A**; Kruskal-Wallis: *χ*^*2*^ = 447.82, *p* = 5.71 × 10^−89^). This is consistent with the hypothesis that transcript abundance is under selection and that strong genetic drift associated with smaller population sizes results in the accumulation of increased deleterious variation in expression.

**Figure 1.**
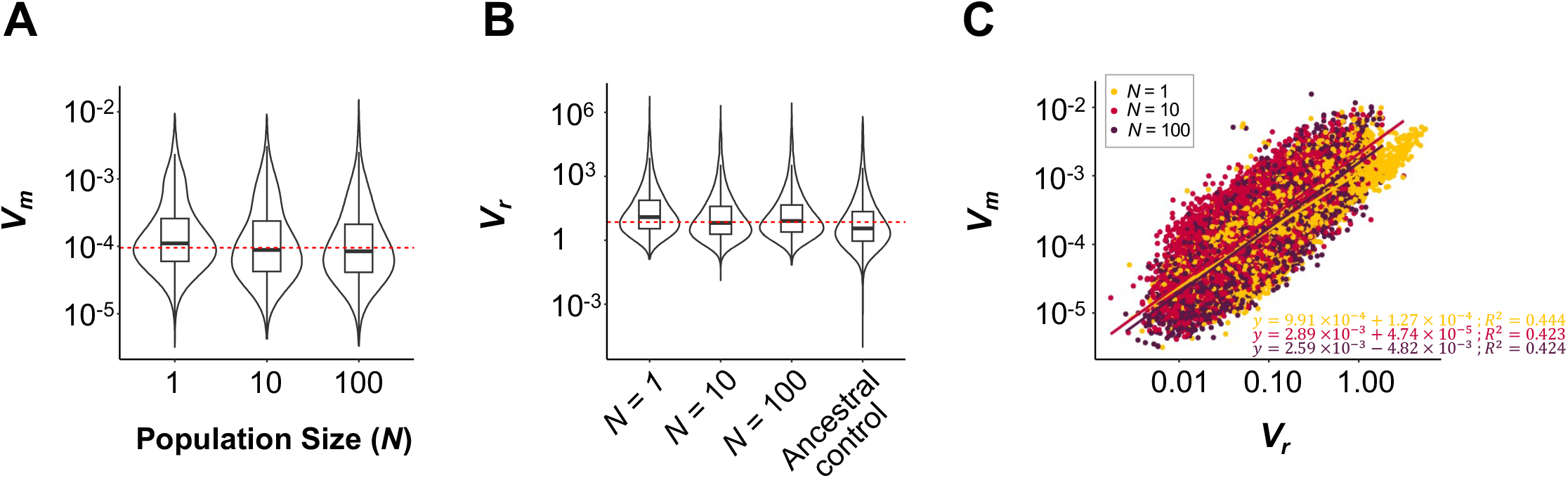
Mutational variance (*V*_*m*_), residual variance (*V*_*r*_), and the relationship between *V*_*m*_ and *V*_*r*_ as a function of population size. *(A) V*_*m*_ shows a negative association with population size across the MA lines. The distributions displayed as violin plots with an internal boxplot indicative of the population median were found to be significantly different (Kruskal-Wallis: *χ*^*2*^= 447.82, *p* = 5.71 × 10^−89^). Median *V*_*m*_ values for the *N* = 1, 10 and 100 lines were 1.11, 0.89, and 0.85 ×10^−4^, respectively. Grand median (9.55 × 10^−5^) for all experimental groups is shown as a dashed red line. *(B)* Median *V*_*r*_ values for *N* = 1, 10, 100 and the ancestral control were 5.89, 3.92, 4.59, and 2.17, respectively. Each of the three MA population size treatments have significantly greater *V*_*r*_than the ancestral control (Dunn’s test: *Z* = 43.13, *p* = 0; *Z* = 22.08, *p* = 4.64 × 10^−108^; *Z* = 29.82, *p* = 2.97 × 10^−195^ for *N* = 1, *N* = 10 and *N* = 100 respectively). The *V*_*r*_ of the *N* = 1 MA lines was significantly greater than in the *N* = 10, and 100 lines (*N* = 1 *vs. N* = 10: Dunn’s test: *Z* = 21.05, *p* = 1.85 × 10^−98^; *N* = 1 *vs. N* = 100: *Z* = 13.31, *p* = 1.28 × 10^−40^). The *V*_*r*_ of *N* = 100 lines was greater than that of the *N* = 10 lines (Dunn’s test: *Z* = 7.74, *p* = 4.96 × 10^− 15^). The grand median (4.41) for all MA lines across the three population size treatments is shown as a dashed red line. *(C)* The correlation between *V*_*m*_ and *V*_*r*_ is show for *N* = 1 (yellow), 10 (red), and 100 (purple) MA lines. The results from linear regression are displayed for each population at the bottom right of the panel.

If relaxation of natural selection results in increased transcriptional dysregulation, the within-line variance in transcript abundance of individual genes (residual variance, *V*_*r*_) would be higher in the *N* = 1 relative to the *N* = 10 and *N* = 100 MA lines. As above, CNVs and TEs were removed prior to the analysis. There was a significant difference in the median TPM *V*_*r*_ values of 5.89, 3.92, and 4.59 for the *N* = 1, 10 and 100 lines, respectively (Kruskal-Wallis: *χ*^*2*^ = 484.97, *p* = 4.89 × 10^−106^). Among the three population size treatments comprising our MA experiment, the *N* = 1 lines were observed to have significantly higher *V*_*r*_ values than the *N* = 10 or 100 MA lines (**Figure 1B**). Extreme relaxation of selection, as in the case of the *N* = 1 MA lines, increases variation in transcript abundance among replicates within lines relative to the larger population size treatments. Furthermore, all of the MA population size treatments resulted in significantly greater *V*_*r*_ compared to the N2 ancestral control which had a median *V*_*r*_ of 2.17 (**Figure 1B**). This suggests that new mutations have rendered transcriptional regulation of some genes less robust and perhaps more sensitive to environmental variation. This contrasts with a previous *C. elegans* study in which the ancestral N2 control was found to have higher *V*_*r*_ than the MA lines (Baer and Denver 2010).

The same features of a gene’s regulatory system that contribute to greater plasticity in expression, whether it is due to a response to some environmental variables or stochasticity in gene regulation, may also result in greater sensitivity of expression to mutations (Meiklejohn and Hartl 2002; Rifkin *et al*. 2005; Landry *et al*. 2007). Genes with greater *V*_*r*_ may therefore also have greater *V*_*m*_, the prediction being a positive correlation between the mutational and residual variance. For this comparison, the *V*_*r*_ was calculated using the same approach as was employed for *V*_*m*_, by generating the log_2_-ratios of MA line replicates to the ancestral control. The median *V*_*r*_ values using this approach were 6.7, 3.7, and 4.8 × 10^−2^ for the *N* =1, 10, and 100 MA lines, respectively. The correlation between *V*_*m*_ and *V*_*r*_ is positive and highly significant (**Figure 1C**; Pearson’s *r* = 0.58, *p* = 0).

### N =1 MA lines exhibit the lowest V_m_/V_r_ ratio

Mutational heritability is defined as the expected heritability of an initially homozygous population following one generation of mutation (Lynch 1988; Houle *et al*. 1996). The ratio of the mutational variance from MA studies to the environmental variance, *V*_*m*_*/V*_*e*_, has been used to estimate the mutational heritability and in the absence of experimental environmental variation, the residual variance, *V*_*r*_, has sometimes been used as a proxy for *V*_*e*_ (Rifkin *et al*. 2005). Following Baer and Denver (2010), we estimated *V*_*r*_ from the variance in gene expression between replicate plates of worms descended from the same MA genotype, wherein each plate housed several thousand L1 larvae. Although *V*_*e*_ should be based on individual variation, we assume that the *V*_*r*_ estimated from the variation among populations (plates) of the same genotype is a function of *V*_*e*_, and can be used to indicate the relative magnitude of *V*_*e*_ between genotypes. There were significant differences between population sizes with respect to *V*_*m*_*/V*_*r*_, with the *N* = 1 lines having significantly lower estimates of *V*_*m*_*/V*_*r*_ relative to the *N* =10 and 100 lines (**Figure 2**). The median estimates of *V*_*m*_*/V*_*r*_ are 1.68, 2.41, and 1.76 ×10^−3^ for the *N* = 1, 10, and 100 lines, respectively (Kruskal-Wallis *χ*^*2*^ = 1225.36, *p* = 8.25 × 10^−267^). This suggests that the mutational heritability in the *N* =1 lines is lower than that of the larger population size treatments. Despite having greater *V*_*m*_, the transcriptional variation generated by new mutations in the *N* = 1 lines may not respond as well to selection in any given environment as the transcriptional variation in the larger populations, either due to greater sensitivity to environmental variation or greater transcriptional noise.

**Figure 2.**
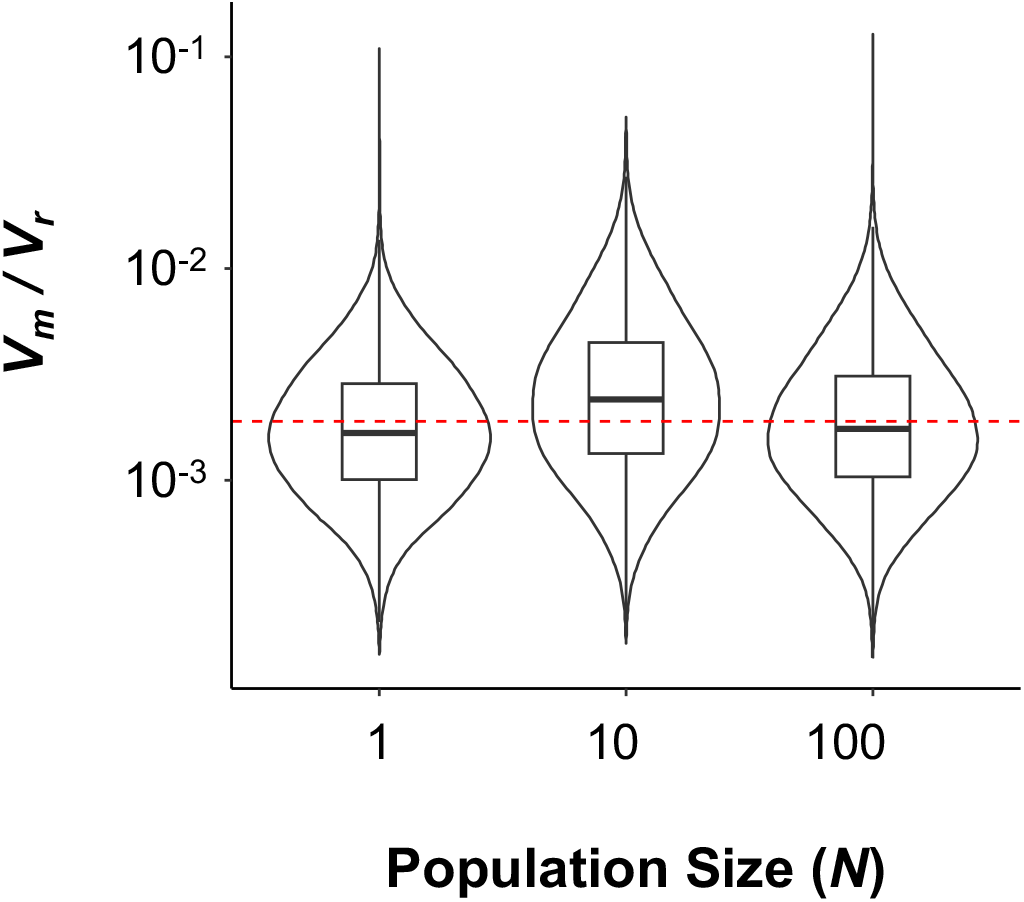
Lower *V*_*m*_ /*V*_*r*_ ratio in the *N* = 1 lines that are under minimal selection efficacy. The *V*_*m*_ /*V*_*r*_ displayed as a violin plot with a median-centered boxplot, was significantly different between the three population size treatments of *N* =1, 10, and 100 (Kruskal-Wallis *χ*^*2*^ = 1225.36, *p* = 8.25 × 10^−267^). The median *V*_*m*_ /*V*_*r*_ for gene expression were 1.68, 2.41, and 1.76 ×10^−3^ for the *N* = 1, 10, and 100 MA lines, respectively.

### Reduced gene expression is a common consequence of mutation accumulation but overexpression may be associated with a greater fitness cost

A moderated T-test (Smyth 2004) was performed to elucidate differential gene expression profiles that developed throughout the course of mutation accumulation by comparing transcript abundance of each MA genotype to that of the N2 ancestral control. We defined significant change in transcript abundance as an increase or decrease of 20% or more, and *p* ≤ 0.01. The average number of genes with significantly increased transcript abundance was 626 ± 121, 367 ± 73 and 326 ± 90 for the *N* = 1, 10 and 100 MA lines, respectively (**Figure 3A**). The average number of genes with significantly decreased transcript abundance was 1088 ± 145, 1251 ± 276, and 1091 ± 354 for *N* = 1, 10, and 100 MA lines, respectively (**Figure 3A**). Irrespective of population size treatment, mutation accumulation disproportionately resulted in reduced transcription of genes (**Figure 3A**; Wilcoxon’s Signed-Ranks *T*_*s*_ = 10, *p* = 2.0 × 10^−8^, *n* = 32 MA lines). Although the number of genes with increased transcription appeared to be greater in the *N* = 1 lines relative to the *N* > 1 lines, the difference was only marginally significant (Kruskal-Wallis *χ*^2^ = 3.78, *p* = 0.051). However, the ratio of up-to down-regulated genes per MA line replicate was significantly different between the *N* = 1 and *N* > 1 lines (Kruskal-Wallis *χ*^2^ = 6.49, *p* = 0.01). This pattern could be suggestive of a greater deleterious fitness effect of gene overexpression and a role of purifying selection in the eradication of such mutations at larger population sizes. Alternatively, the association between genes with increased gene expression and population size could be a consequence of lower fitness or mutations causing transcriptional dysregulation in the *N* = 1 lines. Additionally, we noted that the magnitude and significance of changes in relative transcript abundance were also greater in the *N* =1 MA lines subject to minimal efficiency of selection (**Figure 3B**).

**Figure 3.**
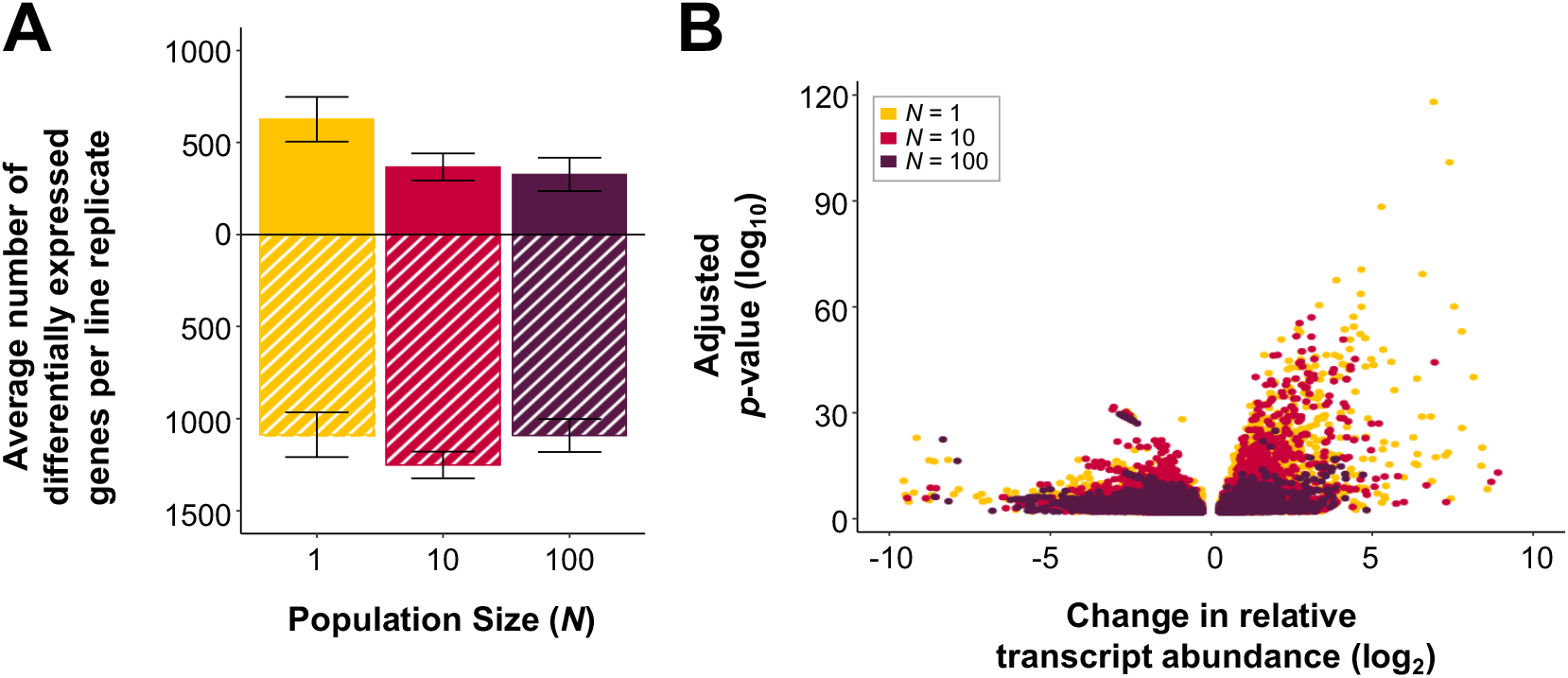
Number of differentially expressed genes and the magnitude of change are increased in lines governed by random genetic drift. MA lines comprising the *N* = 1, 10, 100 population size treatments are displayed in yellow, red, and purple respectively. *(A)* The average number of genes per MA line replicate that have significantly elevated or reduced transcript abundance relative to their N2 ancestor (*p* ≤ 0.01). Only genes that display >20% increase or decrease in transcript abundance are included. Upregulated and downregulated genes are displayed with a solid bar above zero and a hatched bar below zero, respectively. Error bars represent the ± standard error for each population size treatment. *(B)* A volcano plot depicting genes with significantly elevated or reduced transcript abundance relative to the N2 ancestral control. The horizontal axis displays the change in relative transcript abundance as log_2_ of the difference from the ancestral line. The vertical axis represents the log_10_ of adjusted *p*-value for each gene.

### GO enrichment analysis reveals patterns of differential expression between population size treatments

Gene ontology (GO) analyses were performed on differentially expressed genes within each line as identified in the previous section (**Figure 3**). A Fisher’s exact test (*p* ≤ 0.01) was used to determine a significant enrichment with a minimum of 50 genes per GO category. For population sizes of *N* = 1, 10, and 100, we identified significant enrichment in 108, 108, and 85 biological process (BP) categories, 44, 38, and 39 cellular component (CC) categories, and 40, 25, and 23 molecular function (MF) categories, respectively. A complete table of enriched GO terms can be found in **Supplemental Table S2**.

For every GO term enriched within each population size treatment, we further determined the average proportion of genes that were significantly upregulated or downregulated relative to the ancestral state within each experimental line. This analysis was performed on genes meeting significance (*p* ≤ 0.01) and log_2_(± 0.26) thresholds. The four GO categories with the greatest difference in the proportion of upregulated genes between population size treatments were mitochondrial membrane, immune system, response to biotic stimulus and unfolded protein response (**Figure 4**). In all cases, the relative proportion of significantly upregulated genes was greater within the *N* = 1 lines relative to the larger populations (*N* = 10, 100). Additionally, the average log_2_-fold change of immune-related and mitochondrial genes was found to be negatively correlated with MA line fitness (immune-related: Pearson’s *r* = −0.41, *p* = 2.52 × 10^−3^; mitochondrial: Pearson’s *r* = −0.61, *p* = 2.47 × 10^−6^; **Supplemental Figure S2;** Katju *et al*. 2015). The four GO categories with the greatest difference in the proportion of downregulated genes between population size treatments were transcriptional regulation, non-coding RNA processing, chromosome organization and ribosome biogenesis (**Figure 4**). For these categories, the *N* > 1 lines had a greater number of significantly downregulated genes. For comparison, **Figure 4** also includes the proportion of upwardly and downwardly expressed gene for two GO categories that exhibited the least difference between the three population size treatments, namely enzyme inhibitor activity and lysosomes.

**Figure 4.**
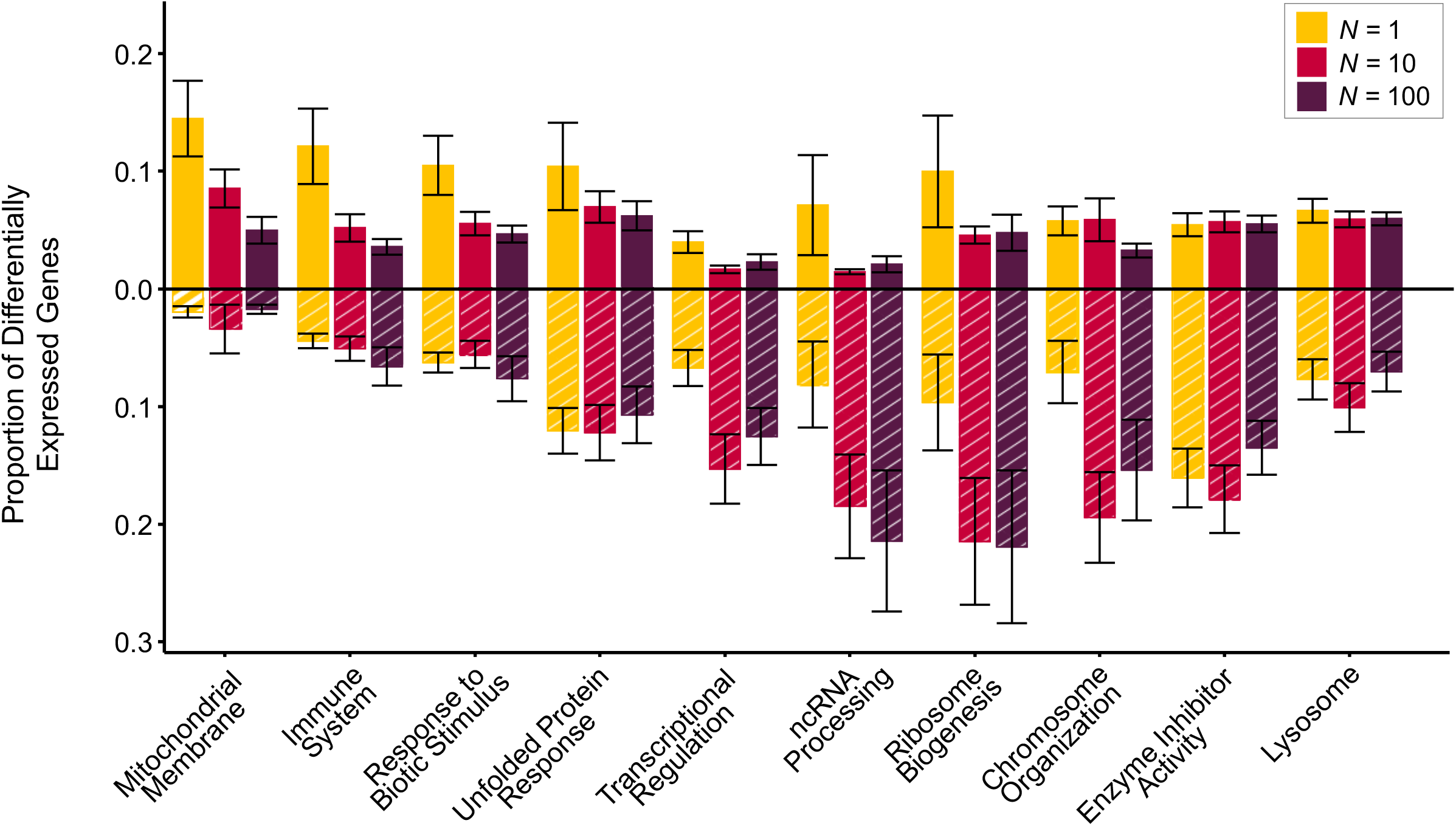
The proportion of upregulated and downregulated genes within specific GO categories highlight greater upregulation in the *N* = 1 lines relative to other population size treatments. The average proportion of significantly upregulated (top, solid bars) and downregulated (bottom, striped bars) genes per line (± standard error) relative to the ancestral N2 control is displayed for select GO categories.

### Histone modification domains influence variation in transcript abundance

Histone modification patterns for *C. elegans* were broadly divided into five domains (Liu *et al*. 2011) associated with (i) broad gene silencing (H3K27me3), (ii) repression in repetitive DNA regions (H3K9me1/me2/me3), (iii) dosage compensation (H3K27me1 and H4K20me1), (iv) activation within promoter regions (H3K4me2, H3K4me3, H4K8ac, H4K16ac, H4tetra-ac, and HTZ-1), and (v) active transcription (H3K79me1/me2/me3 and H3K36me3). We analyzed the variation in transcript abundance between genomic regions associated with different histone modification domains. The average and median Coefficient of Variation (CV) per gene were significantly higher within all population size treatments relative to the ancestral control lines (**Figure 5**). Furthermore, the greatest variation was found between the *N* = 1 lines (**Figure 5**). The median CV for chromatin domains associated with repression for populations *N* = 1, 10, 100 and the ancestral control were 1.7, 1.42, 1.55, 1.03 × 10^−1^ (Kruskal-Wallis *χ*^*2*^ = 628.95, *p* = 5.32 × 10^−136^). The median CV for genes associated with silencing in repetitive DNA regions was 1.34, 1.11, 1.24, 0.95 × 10^−1^ (Kruskal-Wallis *χ*^*2*^ = 121.71, *p* = 3.31 × 10^−26^) for populations *N* = 1, 10, 100 and the ancestral control, respectively. For genes associated with dosage compensation, median CV values were 1.25, 0.99, 1.04, 0.89 × 10^−1^ (Kruskal-Wallis *χ*^*2*^ = 228.66, *p* = 2.70 × 10^−49^) for *N* = 1, 10, 100 lines and the ancestral control, respectively. Variation in transcript abundance of genes associated with transcriptional activation was notably lower than their repressed counterparts, with a median CV of 1.18, 0.97, 1.07, 0.82 × 10^−1^ (Kruskal-Wallis *χ*^*2*^ = 686.49, *p* = 1.78 × 10^−148^) for *N* = 1, 10, 100 lines and ancestral control, respectively. Finally, the median CV associated with histone modification denoting constitutively expressed genes was 1.15, 0.92, 0.98, 0.87 × 10^−1^ (Kruskal-Wallis *χ*^*2*^ = 140.39, *p* = 3.11 × 10^−30^) for *N* = 1, 10, 100 lines and ancestral control, respectively.

**Figure 5.**
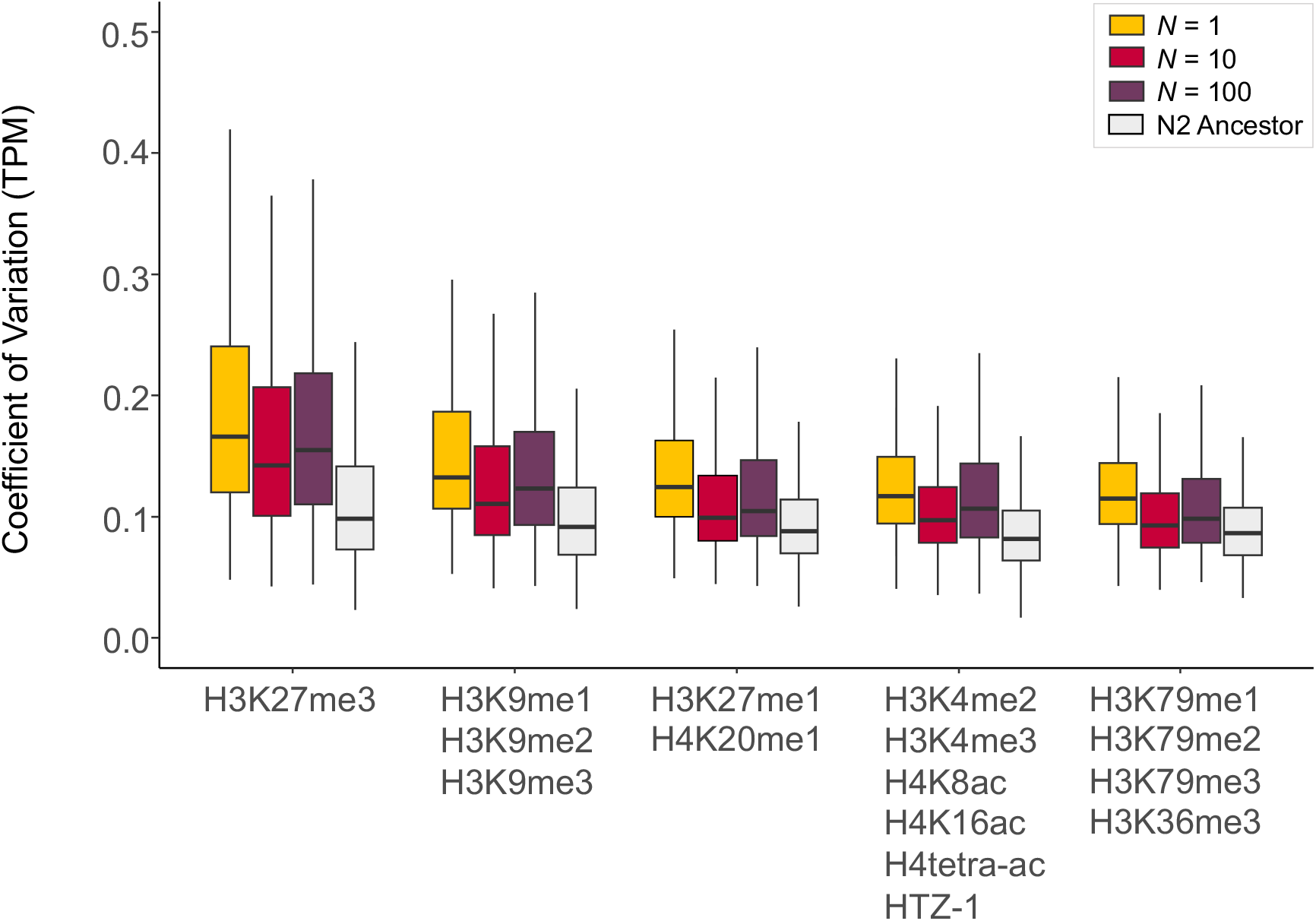
The Coefficient of Variation (CV) for genes associated with specific chromatin states are significantly higher in lines governed by extreme genetic drift (*N* = 1). The boxplot represents the distribution of CV for genes located in genome regions associated with the histone modifications indicated on the x-axis. Significant differences were found between population sizes (*N* = 1, 10, and 100 individuals) for all chromatin states (Kruskal-Wallis: H3K27me3, *χ*^*2*^ = 154.35, *p* = 3.04 × 10^−34^; H3K9me1/me2/me3, *χ*^*2*^ = 35.47, *p* = 1.99 × 10^−8^; H3K27me1, H4K20me1, *χ*^*2*^ = 102.74, *p* = 4.91 × 10^−23^; H3K4me2/me3, H4K8ac, H4K16ac, H4K16ac, H4Ktetra-ac, HTZ-1, *χ*^*2*^ = 186.93, *p* = 2.57 × 10^−41^; H3K79me1/me2/me3, H3K36me3, *χ*^*2*^ = 72.58, *p* = 1.74 × 10^−16^). The differences in CV for individual genes within the three different population size treatments are highly significant between chromatin states (Kruskal-Wallis: *N* = 1, *χ*^*2*^ = 648.00, *p* = 6.31 × 10^−139^; *N* = 10, *χ*^*2*^ = 628.78, *p* = 9.15 × 10^−135^; *N* = 100, *χ*^*2*^ = 635.84, *p* = 2.70 × 10^−136^).

## DISCUSSION

Mutation accumulation (MA) studies, originally conceived in the 1960s to indirectly estimate mutational parameters including the deleterious mutation rate from fitness-related phenotypic data, have emerged as a powerful framework to directly determine the rate and spectrum of spontaneous mutations as well as their transcriptional consequences. The last decade has witnessed a burgeoning of MA experiments that have served to increase species representation, thereby offering insights into mutational parameters in emerging model species of prokaryotes as well as new eukaryotic taxa such as algae, amoeba, ciliates, crustaceans, and mammals (reviewed in Katju and Bergthorsson 2019). Spontaneous MA experiments provide a powerful framework to assess global transcriptional divergence due to the input of new genetic variation via mutation. Yet, to date, this approach remains surprisingly underutilized. This study using *C. elegans* represents the first attempt to analyze global transcriptional divergence in MA lines using (i) next-generation sequencing technology (RNA-Seq) in conjunction with (ii) a modified MA design comprising differing population size treatments to additionally discern the effects of mutation and selection on transcriptional divergence.

Comparisons of transcriptional divergence within natural isolates of species (Hutter *et al*. 2008; Hodgins-Davis *et al*. 2015), between species (Lemos *et al*. 2005; Rifkin *et al*. 2005; Hodgins-Davis *et al*. 2015) and MA experiments (Denver *et al*. 2005; Rifkin *et al*. 2005; McGuigan *et al*. 2014a; Hodgins-Davis *et al*. 2015; Huang *et al*. 2016) suggest that gene expression is under strong stabilizing selection. Stabilizing selection on gene expression predicts that mutational variance (*V*_*m*_) in transcript abundance is greatest where the effects of genetic drift are predicted to be highest, such as in our *N* = 1 MA lines. Our results are consistent with this prediction in that the *N* = 1 MA lines exhibit significantly greater mutational variation in transcript abundance relative to larger population size treatments (*N* = 10, 100 MA lines). We have previously reported that regulation of transposable element transcription is also under selection in these MA lines and that TE transcription increased with time, especially in the *N* = 1 lines (Bergthorsson *et al*. 2020). Furthermore, gene copy-number variants that substantially increased transcript abundance were less likely to accumulate in larger populations relative to the MA lines subjected to the most extreme population bottleneck, which demonstrated the importance of expression of gene duplicates for fitness (Konrad *et al*. 2018). In the analyses presented here, TEs and CNVs were filtered out. The number of mutations per generation that accumulated in these MA lines was not significantly different between the three population size treatments (Konrad *et al*. 2019), but it nevertheless appears that mutations altering change gene expression are associated with reduced fitness, and have been selected against in MA lines with larger population bottlenecks.

The residual variance, *V*_*r*_, reflecting the within-line variation in gene expression was significantly greater in the MA lines than in the N2 ancestral control. Furthermore, *V*_*r*_ was significantly greater in the *N* = 1 MA lines relative to their *N* > 1 counterparts. As with the mutational variance, this suggests that mutations resulting in increased sensitivity to environmental and transcriptional noise due to dysregulation of gene expression are accumulating to the greatest degree under conditions of extreme genetic drift, and to a lesser degree in MA lines with larger population bottlenecks (and greater efficiency of selection). This increase in the *V*_*r*_ for transcription in our MA lines contrasts with a previous *C. elegans* study using DNA microarrays in which the ancestral N2 control was found to have higher *V*_*r*_ than the MA lines (Baer and Denver 2010). The observation that an increase in *V*_*r*_ largely mirrors the pattern for *V*_*m*_ suggests that the two may be related and perhaps stem from the same mechanistic causes.

There was a strong positive correlation between the per gene *V*_*m*_ and *V*_*r*_. This correlation is in accord with results from other experimental systems such as *S. cerevisiae* and *D. melanogaster* where mutational variation in gene expression was determined to be positively associated with environmental variation and gene expression noise (Rifkin *et al*. 2005; Landry *et al*. 2007; Huang *et al*. 2016). The canalization of gene expression, which renders patterns of gene expression robust to the effects of environmental variation, also appears to buffer the perturbing effects of mutations by similar means (Wagner *et al*. 1997; Meiklejohn and Hartl 2002; Rifkin *et al*. 2005; Landry *et al*. 2007; Baer and Denver 2010; Huang *et al*. 2016).

While the mutational variance (*V*_*m*_) in transcript abundance was negatively correlated with population size, the *V*_*m*_/*V*_*r*_ ratio for transcript abundance unexpectedly revealed the opposite relationship (smaller in the *N =* 1 lines than in the *N* > 1 MA lines). This was owing to a disproportionate increase in residual variance, *V*_*r*_, in the *N* = 1 lines relative to the *N* > 1 MA lines. Decanalization is often portrayed as increasing evolvability by revealing cryptic genetic variation (Wagner *et al*. 1997; Gibson and Dworkin 2004; Flatt 2005; Baer 2008). Conversely, if canalization makes phenotypes robust to genetic and environmental perturbation by the same mechanisms, decanalization may also decouple the genotype and phenotype by increasing microenvironmental sensitivity and noise in gene expression to such a degree that natural selection becomes less effective in the short-term. However, the fact that higher *V*_*r*_ and a lower *V*_*m*_/*V*_*r*_ ratio (predicted to be correlated with mutational heritability) are observed in the *N* = 1 lines suggests that decanalization of a large number of genes is itself often deleterious, and would be selected against under realistic conditions in natural populations.

We investigated the influence of chromatin domains on the variation in gene expression utilizing five broad domain categories that have been previously defined (Liu *et al*. 2011). The median CV of transcript abundance in the ancestral control of the MA lines was very similar between genes in different domains. However, following mutation accumulation, the median CV increased for genes in all five domains across all population size treatments. The greatest and smallest increase in median CV was for genes associated with broad gene silencing and active transcription, respectively. Within each domain category, the *N* = 1 lines always exhibited the greatest increase in median CV for gene expression. For example, in repressed domains, the median CV in the *N* = 1 lines increased by 65% compared to a 37% and 50% increase in the *N* = 10 and *N* = 100 lines, respectively. In constitutively active domains, the median CV in the *N* = 1 lines increased by 32% compared to a 13% and 6% increase in the *N* > 1 lines (*N =* 10 and *N =* 100). Genes in domains associated with silencing of repeats, dosage compensation and activated transcription fell between these extremes. It appears that the regulation of genes in repressed domains is most sensitive to genetic perturbation from the accumulation of mutations.

Mutations can alter gene expression, thereby resulting in phenotypic divergence from the wildtype. In terms of transcript abundance, this dysregulation of gene expression following mutation can manifest in two modes, namely (i) underexpression, or (ii) overexpression. Both these deviations from wildtype expression levels have the potential to alter phenotypes in a detrimental or beneficial manner, depending on the environmental context. There are inherent challenges in our ability to predict if, and how, a particular new mutation will influence gene expression, especially if it is *trans*-acting (Lutz *et al*. 2019; Hill *et al*. 2020). Nevertheless, we sought to investigate if spontaneous mutations confer any broad, general patterns with respect to expression dysregulation under minimal and increasing intensity of selection. More genes were significantly underexpressed than overexpressed after mutation accumulation across all population size treatments. However, the ratio of significantly overexpressed to underexpressed genes was greater in the *N* = 1 MA lines. This pattern could be suggestive of a greater deleterious fitness effect of gene overexpression and a role of purifying selection in the eradication of such mutations at larger population sizes. Alternatively, the association between genes with increased gene expression and population size could be a consequence of the, on average, lower fitness of the *N* = 1 lines (Katju *et al*. 2015, 2018). Some alterations in gene expression could, for example, be compensating for mutations causing transcriptional dysregulation of other genes.

An examination of gene expression profiles offers a direct approach to investigate whether genes are equally malleable in their ability to diverge at the transcriptional level. Genes associated with the mitochondrial membrane and immune system exhibited the greatest average increase in expression over ancestral levels in the MA lines. The vigorous upregulation of immune response genes was especially pronounced in low fitness (*N* = 1) lines **(Figure 5)** despite the absence of a pathogenic challenge. While the standard *C. elegans* practice of feeding nematodes live OP50 (*E. coli*) has been documented to decrease lifespan compared to nematodes fed dead bacteria in a model of slow pathogenesis (Battisti *et al*. 2016; Darby 2005), the L1 larvae sequenced for this experiment were not exposed to food, living or dead, following synchronization. Furthermore, while these *N* =1 lines possessed the greatest degree of upregulation in downstream inflammatory response genes, we did not observe any significant expression changes in key regulatory genes upregulated directly in response to bacterial challenge, such as *pmk-1/pmk-2, sek-1*, and *nsy-1* (Kim *et al*. 2002). However, infection response genes in *C. elegans* share significant overlap with stress response pathways and can be triggered by cellular stress or damage (Matzinger 1994). Because immune effectors increase ER stress and increased expression in the unfolded protein response (UPR) pathway, another category predominantly enriched in the *N* = 1 lines (**Figure 5**), we turned focus on to the *ire-1/xbp-1* pathway. This pathway has been documented to confer host protection via alleviation of ER stress resulting from immune system activation (Calfon *et al*. 2002; Shapira *et al*. 2006; Richardson *et al*. 2010). Only a single line, 1N, was found to possess significant upregulation of genes within this network, presumptively providing protection against the detrimental effect of increased inflammation. Thus, the negative fitness effects brought about by voracious activation of immune pathways might be coupled with increased ER stress and a lack of concurrent increases in pathways involved in managing such stresses.

Previous analyses into the fitness effects of mutations in lines of *C. elegans* MA lines with population bottlenecks of *N* = 1, 10 and 100 found no significant fitness decline in the *N* = 10, and 100 lines (Katju *et al*. 2015, 2018). Furthermore, wholegenome sequencing found no differences in the accumulation of base substitutions and small indels among the MA lines of different population sizes (Konrad *et al*. 2019). The molecular differences that we have found between MA lines that were maintained under different efficiency of selection (different population sizes) are primarily associated with expression: increased expression and increased variation in expression of transposable elements (Bergthorsson *et al*. 2020), greater expression of duplicated genes (Konrad *et al*. 2018), and greater variation of expression in other genes. It has been noted that the number of genes with significant differential expression in MA lines is at least an order of magnitude greater than the number of mutations per genome (Gibson 2005), which is confirmed in this study. These results suggest that mutations that cause large-scale changes in gene expression are more deleterious than those that alter gene expression to a lesser degree. This is also in accord with our previous results regarding TEs and expression of gene duplicates. However, some of the changes in gene expression in low fitness lines may be attempts to ameliorate the effects of prior deleterious mutations. For example, the negative correlation between fitness and increased transcription of immune/stress/unfolded protein response genes may be examples of compensatory response in gene expression rather than contributing directly to fitness decline. Our study extends previous findings regarding balancing selection on gene expression. Furthermore, we also observed significant expression divergence in MA lines that did not exhibit significant fitness loss. However, the population sizes used in these experiments are too small to conclude that this expression divergence is neutral.

## ACKNOWLEDGMENTS

This research was partially supported by National Science Foundation grant MCB1565844 to V.K. U.B. and V.K. were additionally supported by start-up funds from the Department of Veterinary Integrative Biosciences, College of Veterinary Medicine and Biomedical Sciences at Texas A&M University.

## AUTHOR CONTRIBUTIONS

TCD, Acquisition of data, Analysis and interpretation of data, Drafting or revising the article; AK, Acquisition of data, Analysis and interpretation of data; UB and VK, Conception and design, Analysis and interpretation of data, Drafting or revising the article.

## COMPETING INTERESTS

The authors declare that no competing interests exist.

## ADDITIONAL FILES

### Supplementary files

- Supplementary file 1. Supplemental Tables S1 and S2.
- Supplementary file 2: Supplemental Figure S1
- Supplementary file 3: Supplemental Figure S1

### Major Datasets

The sequence data generated in this study is available in Bioproject PRJNA448413 with accession code SRP145329 (Bioproject link: https://www.ncbi.nlm.nih.gov/bioproject/PRJNA448413).

## Notes

https://www.ncbi.nlm.nih.gov/bioproject/PRJNA448413

### Competing Interest Statement

The authors have declared no competing interest.

