## Supplemental Figure S1 for "Global Gene Expression Divergence in Spontaneous Mutation Accumulation Lines of *Caenorhabditis elegans* under Varying Efficiency of Selection"

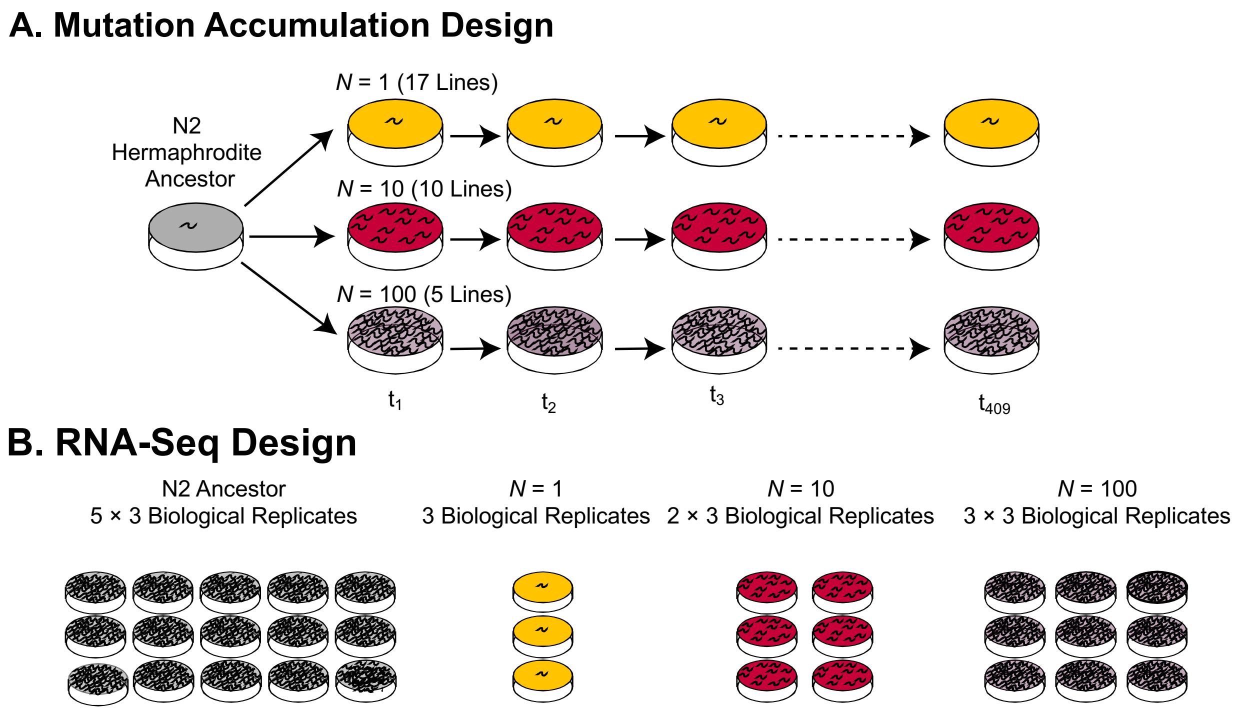


**Supplemental Figure S1. Design of mutation accumulation (MA) experiment and RNA-Seq.** *(A)* Independent MA lines descended from a single N2 hermaphrodite ancestor were maintained at three population sizes of $N = 1$ (17 lines), $N = 10$ (10 lines), and $N = 100$ (5 lines) over 409 generations. Additional descendants of the same N2 ancestor were expanded for two generations and cryopreserved as ancestral, pre-MA controls. The maintenance of lines at varying $N$ enables manipulation of the strength of selection. Following several hundred generations, the $N = 1$ lines were expected to have independently accumulated mutations under a regime of strong genetic drift and minimal selection, leading to a mean decline in fitness relative to the ancestral control and increased among-line variance. The larger population size treatments ($N = 10$ and 100 worms) were subject to greater selection intensity. These larger population size treatments were expected to accumulate mutations whose fates were determined by the fitness effects of the mutation as well as the strength of natural selection operating in that particular genetic background. *(B)* Experimental design for RNA-Seq at the L1 larval stage. One, two and three individuals were isolated from each MA lines of size $N =$ 1, 10, and 100 individuals respectively. Additionally, five individuals from the ancestral pre-MA control were isolated for sequencing. For each of these 57 individuals, three pre-adult L4 offspring were isolated from the $F_1$ generation to establish three biological replicates yielding a total of 171 lines for L1 tissue collection and RNA-Seq.
