## Supplemental Figure S2 for "Global Gene Expression Divergence in Spontaneous Mutation Accumulation Lines of *Caenorhabditis elegans* under Varying Efficiency of Selection"

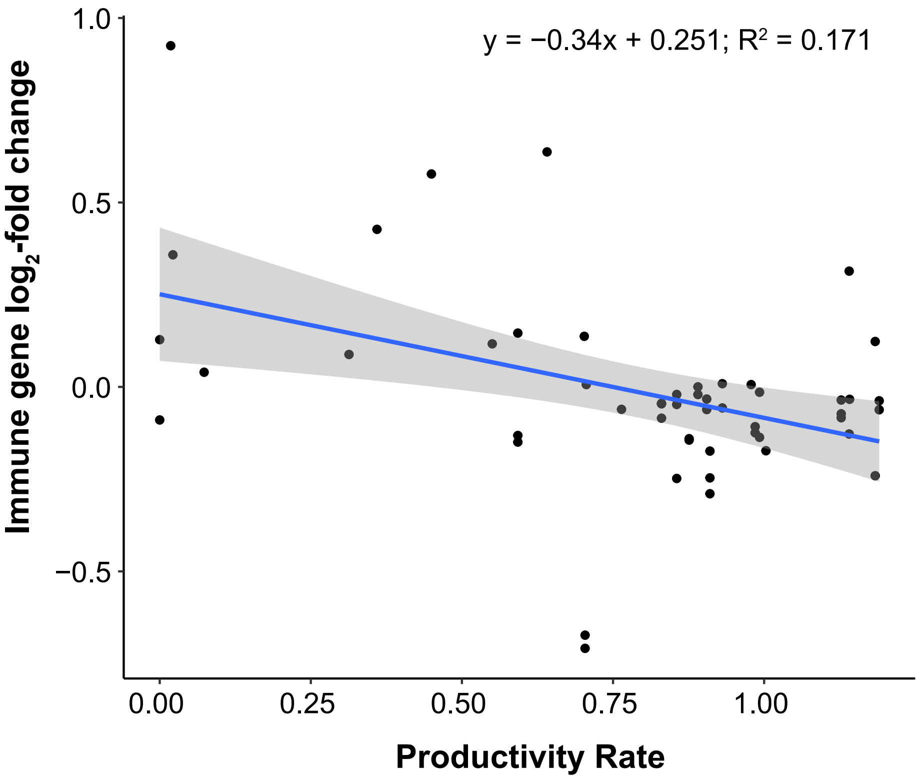


**Supplemental Figure S2. Immune-related gene expression is negatively correlated with productivity rates of experimental lines.** The average $\log_2$-fold change for immune-related genes was calculated and plotted against the mean productivity rate for each MA line. The regression line (blue, standard error in grey) equation is displayed at the top of the graph. The regression analysis identified a significant relationship ($p = 0.002$) between upregulation of immune genes and diminished fitness.
